# Cooperate-and-radiate co-evolution between ants and plants

**DOI:** 10.1101/306787

**Authors:** Katrina M. Kaur, Pierre-Jean G. Malé, Erik Spence, Crisanto Gomez, Megan E. Frederickson

## Abstract

Mutualisms may be “key innovations” that spur diversification in one partner lineage, but no study has evaluated whether mutualism accelerates diversification in both interacting lineages. Recent research suggests that plants that attract ant mutualists for defense or seed dispersal have higher diversification rates than non-ant associated plant lineages. We ask whether the reciprocal is true: does the ecological interaction between ants and plants also accelerate diversification in ants? In other words, do ants and plants cooperate-and-radiate? We used a novel text-mining approach to determine which ant species associate with plants in defensive or seed dispersal mutualisms. We investigated patterns of trait evolution and lineage diversification using phylogenetic comparative methods on a large, recent species-level ant phylogeny. We found that ants that associate mutualistically with plants have elevated diversification rates compared to non-mutualistic ants, suggesting that ants and plants cooperate-and-radiate.

## Introduction

How have species interactions contributed to the diversification of life on Earth? Ehrlich and Raven^1^ famously proposed escape-and-radiate co-evolution as an engine of plant and insect diversification. Here we present evidence that “cooperate-and-radiate” co-evolution is also an important diversifying force. Recent studies have linked mutualism evolution to accelerated lineage diversification^2–4^, suggesting that mutualism either buffers lineages against extinction, promotes speciation as lineages enter new adaptive zones, or both^5^. Building on previous research showing that partnering with ants enhances plant diversification^3,4^, we ask if interacting mutualistically with plants enhances ant diversification.

Mutualism theory generally predicts the opposite; mutualism is expected to hinder diversification^6,7^ because the interdependence of partners, conflicts of interest between them, or the invasion of selfish “cheaters” should make mutualistic lineages vulnerable to extinction^8–10^. However, the available phylogenetic evidence strongly suggests that mutualisms often persist over long periods of evolutionary time, and may help lineages flourish^8^. Furthermore, recent studies show that mutualism can expand a lineage’s realized niche^11,12^, potentially creating ecological opportunity.

Ant-plant interactions are classic examples of mutualism that have evolved numerous times in both partner lineages, but a relationship between ant-plant mutualisms and enhanced diversification has been tested for only in plants^3^’^4^’^13^. Ant “bodyguards” visit extrafloral nectaries (EFNs) on plants or nest in specialized plant cavities (domatia) and protect plants against herbivores or other enemies^14^. Ants also disperse seeds that bear lipid-rich elaiosomes^3^. The evolution of elaiosomes^3^ and EFNs^4^, but not domatia^13^, enhances plant diversification, providing one possible explanation for the rapid radiation of angiosperms famously referred to as “Darwin’s abominable mystery”^15^. Ants can reduce the negative effects of seed predators or herbivores on plant populations, potentially reducing extinction risk^4,16^, or they may help plants colonize new sites, promoting speciation^3,16^, although it is worth emphasizing that ants move seeds only short distances^17^. Since ant and angiosperm radiations are broadly contemporaneous, having diversified during the Late Cretaceous^18–20^, they may have ‘cooperated-and-radiated’; on the ant side, the evolution of ant-plant interactions may have made new niches available in the form of plant-derived food or nest sites, resulting in expanded ranges or decreased extinction risk.

To investigate cooperate-and-radiate co-evolution, we automated the compilation of trait data from the primary literature. Inspired by recent studies that have leveraged advances in bioinformatic pipelines^21^ and machine reasoning^22^ to characterize, for example, protein-protein interaction networks^23^, we took an innovative text-mining approach to compile ant-plant interaction data from the abstracts and titles of over 89,000 ant-related publications. We groundtruthed the results by manually checking a subset of abstracts for false positives and evaluated how the number of unique plant-ant species identified by our text-mining algorithm changed as the algorithm sampled more abstracts. We analyzed our trait data in conjuction with a species-level ant phylogeny using both Binary State Speciation and Extinction (BiSSE)^24^ and Bayesian Analysis of Macroevolutionary Mixtures (BAMM)^25^ models to assess the effect of mutualism on ant diversification. Thus, we evaluate for the first time whether the evolution of mutualism leads to reciprocal diversification in ants and plants through cooperate-and-radiate co-evolution.

## Results

Text-mining generated a wealth of trait data (Figure 1, Figure S1). This method outputted trait data for 3342 ant species in 265 ant genera, representing 23% and 73%, respectively, of all currently recognized ant species and genera, globally. The text-mining extracted these data from approximately 15,000 abstracts, with the number of unique species increasing as the number of abstracts sampled increased (Figure 2). The species accumulation curves suggest that our lists of ant species that nest in domatia or consume plant food bodies are relatively complete, but that text-mining an even a larger sample of abstracts from the primary literature would identify many more ant species that disperse seeds or visit EFNs, as well as many more non-mutualistic ant species (Figure 2).

**Figure 1.**
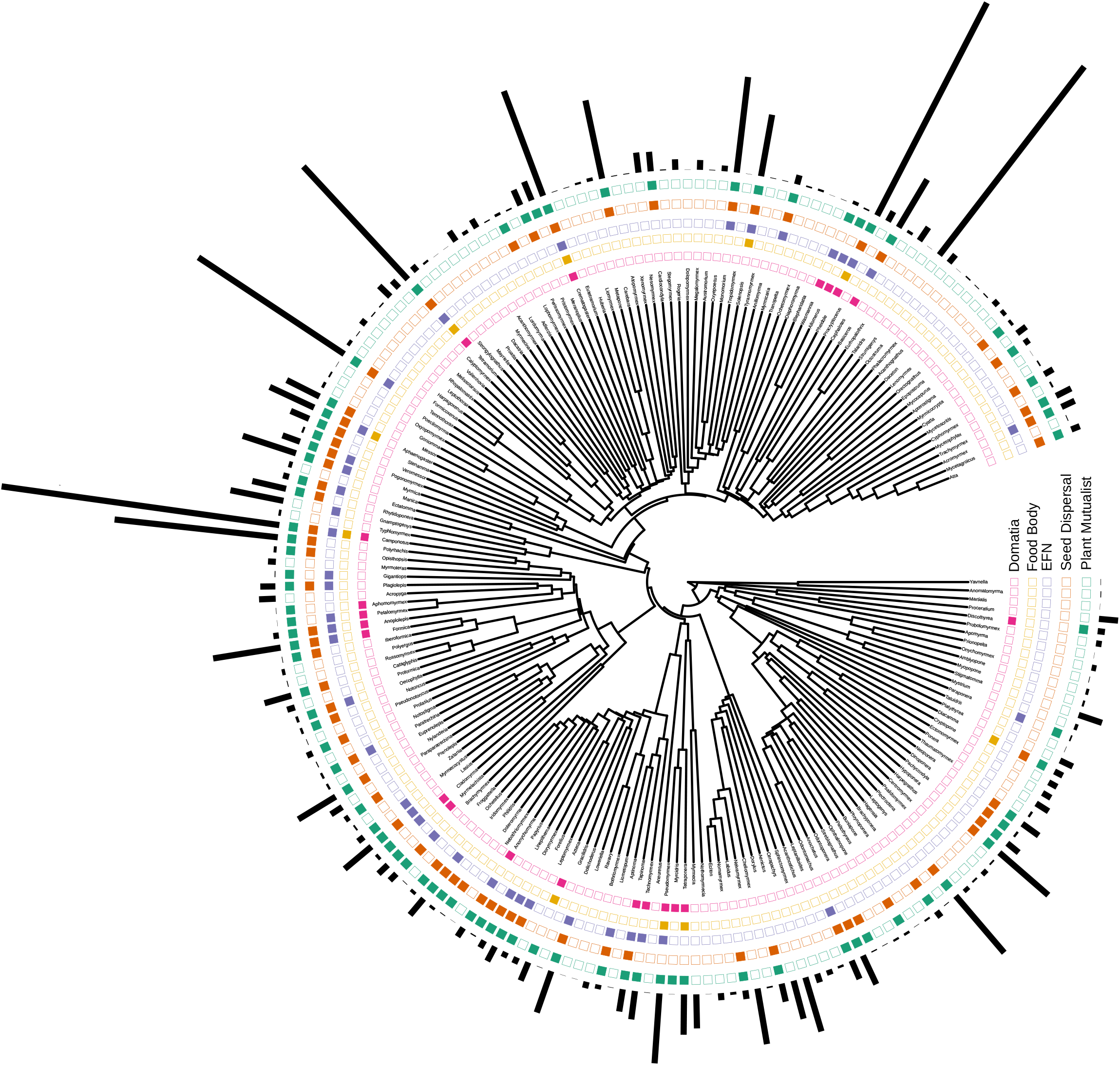
Visualization of trait data, showing which genera (N = 199) contain species that nest in domatia (pink), consume food bodies (yellow), visit EFNs (purple), disperse seeds (orange), or engage in any plant mutualism (combination of all traits) (green). Black bars show the number of species in each genus. The full species-level phylogeny and trait data used in analyses is in the Supplementary Information.

**Figure 2.**
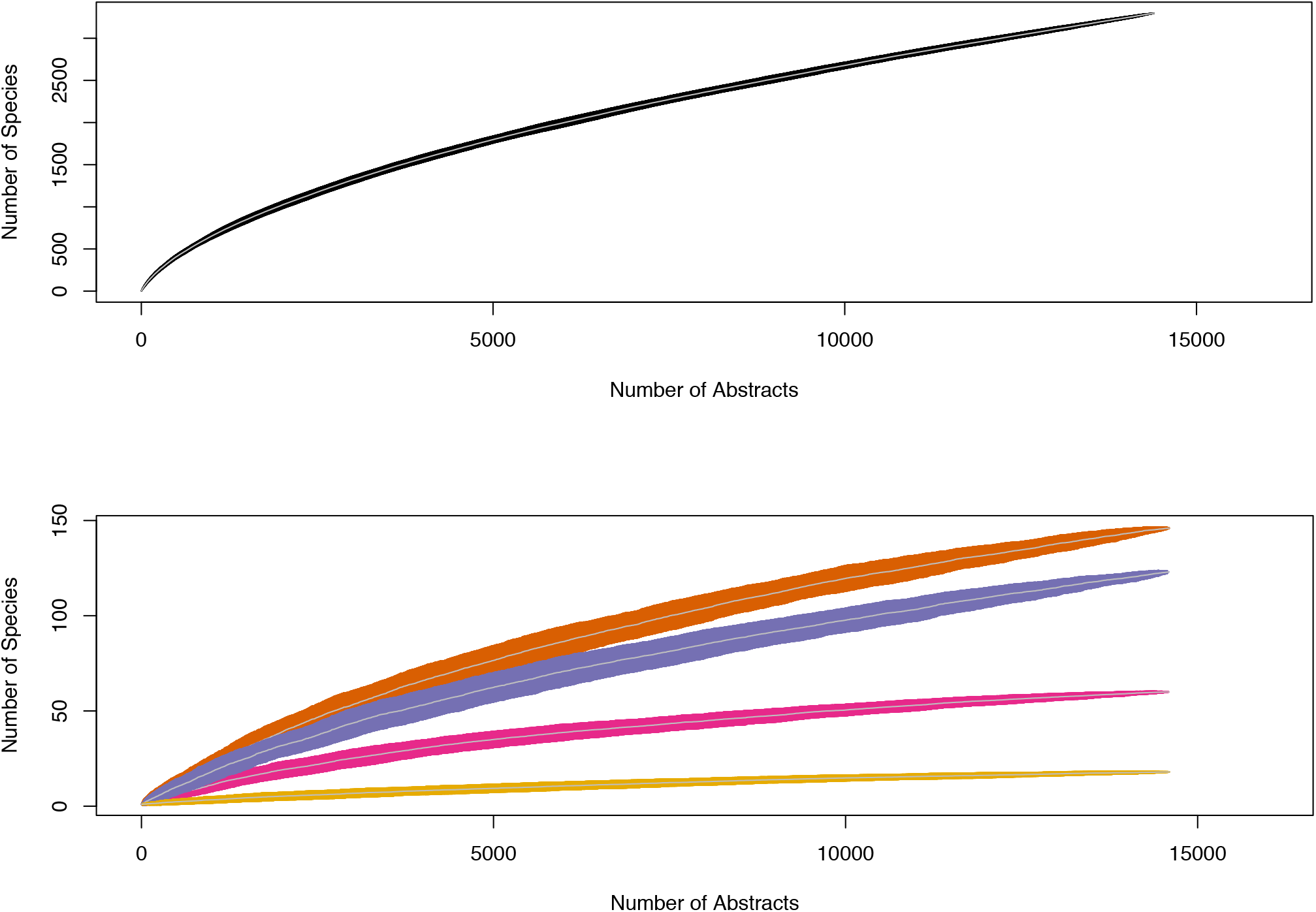
Species accumulation curves showing how the number of unique ant species increased as more abstracts were sampled for both ants that do not interact mutualistically with plants (top panel) and ants that disperse seeds (orange), visit EFNs (purple), nest in domatia (pink), and consume food bodies (yellow), (bottom panel). Nearly 15,000 abstracts contained ant names. Grey lines are means and black or colored regions are standard deviations calculated from sub-sampling abstracts 100 times at each x-axis value.

We paired this approach with a more traditional method of manually assembling a list of seed-dispersing ants by reading the primary literature (183 articles). Our automated approach performed relatively well in comparison; 85 seed-dispersing ant species in 28 genera occurred in both our text-mining results and this hand-compiled dataset. In total, the text-mining identified 129 ant species in 39 genera as seed dispersers, compared to 268 ant species in 60 genera in the hand-compiled dataset, meaning that 44 ant species were in the text-mining results only, and 183 species were in the hand-compiled dataset only. Of the 183 seed-dispersing ant species in the hand-compiled dataset but not in our text-mining output, 149 species were described in 92 papers that were not included among our 89,000 abstracts, suggesting that the largest improvements to our text-mined trait dataset would come from having access to more abstracts, rather than from a better text-mining algorithm. The text-mining method was not immune to error, but the number of false positives declined rapidly as the number of abstracts in which ant names and trait terms co-occurred increased. We found no false positives when an ant species name co-occurred with one or more trait terms in 4 or 5 abstracts (Figure 3).

**Figure 3.**
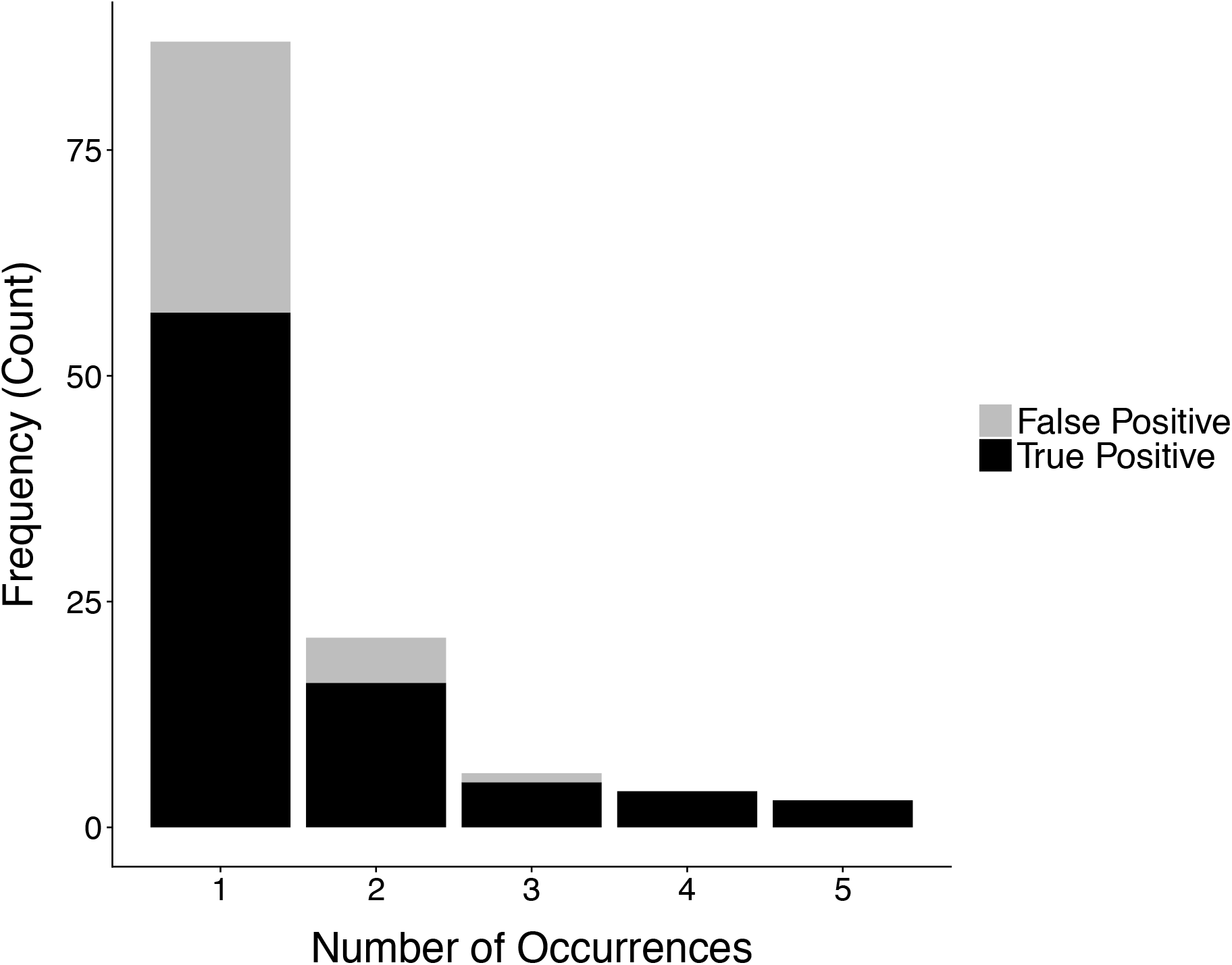
When an ant species name appeared with a trait term in five or fewer abstracts, abstracts were manually scored to check for false positives. No false positives were detected when ant species name and trait terms co-occurred in at least 4 abstracts.

Seed-dispersing and EFN-visiting ants were most commonly identified by text-mining, followed by domatia-nesting ants; relatively few ant species that consume food bodies were found by text-mining. Specifically, we identified 309 ant species in 77 genera that disperse seeds (including taxa in the hand-compiled dataset) and 3030 ant species in 261 ant genera that do not; 122 ant species in 37 genera that visit EFNs and 3182 ant species in 261 ant genera that do not; 58 ant species in 22 genera that nest in domatia and 3246 ant species in 262 genera that do not; 16 ant species in 8 genera that consume plant food bodies (other than elaiosomes) and 3288 species in 265 genera that do not. In all, we identified 432 ant species in 84 genera that are plant mutualists, and 2910 ant species in 256 genera that are not; subsequent analyses were conducted on this combined category.

After pruning the phylogeny to match the trait data, the tree was comprised of 795 species in 199 genera (Figure 1, Figure S1). Unlike other studies of trait-dependent diversification, we did not include species for which we had no trait information (i.e., ant species that did not appear in our text-mined abstracts). Thus, the 795 species used in the BiSSE and BAMM analyses were species for which the text-mining determined the ant species is likely a plant mutualist (432 species), as well as species for which the text-mining determined the ant species is likely not a plant mutualist (363 species); the latter ant species names appeared in at least one of our ~89,000 abstracts but were never found together with any plant mutualist terms (Table S2). Both BiSSE and BAMM models found that mutualistic ant lineages diversify faster than non-mutualistic ant lineages (Table 1, Figure 4). In the BiSSE model, there was no overlap between plant mutualists and non-mutualists in their 95% credible intervals for speciation, extinction, and transition rates, meaning these differences are statistically significant (Table S3). Both BiSSE and BAMM analyses reported higher speciation and higher extinction rates when ants evolved mutualisms with plants; however, overall diversification rates were higher in mutualistic ants, despite elevated extinction rates, because of the even greater difference in speciation rates between mutualistic and non-mutualistic ants.

**Figure 4.**
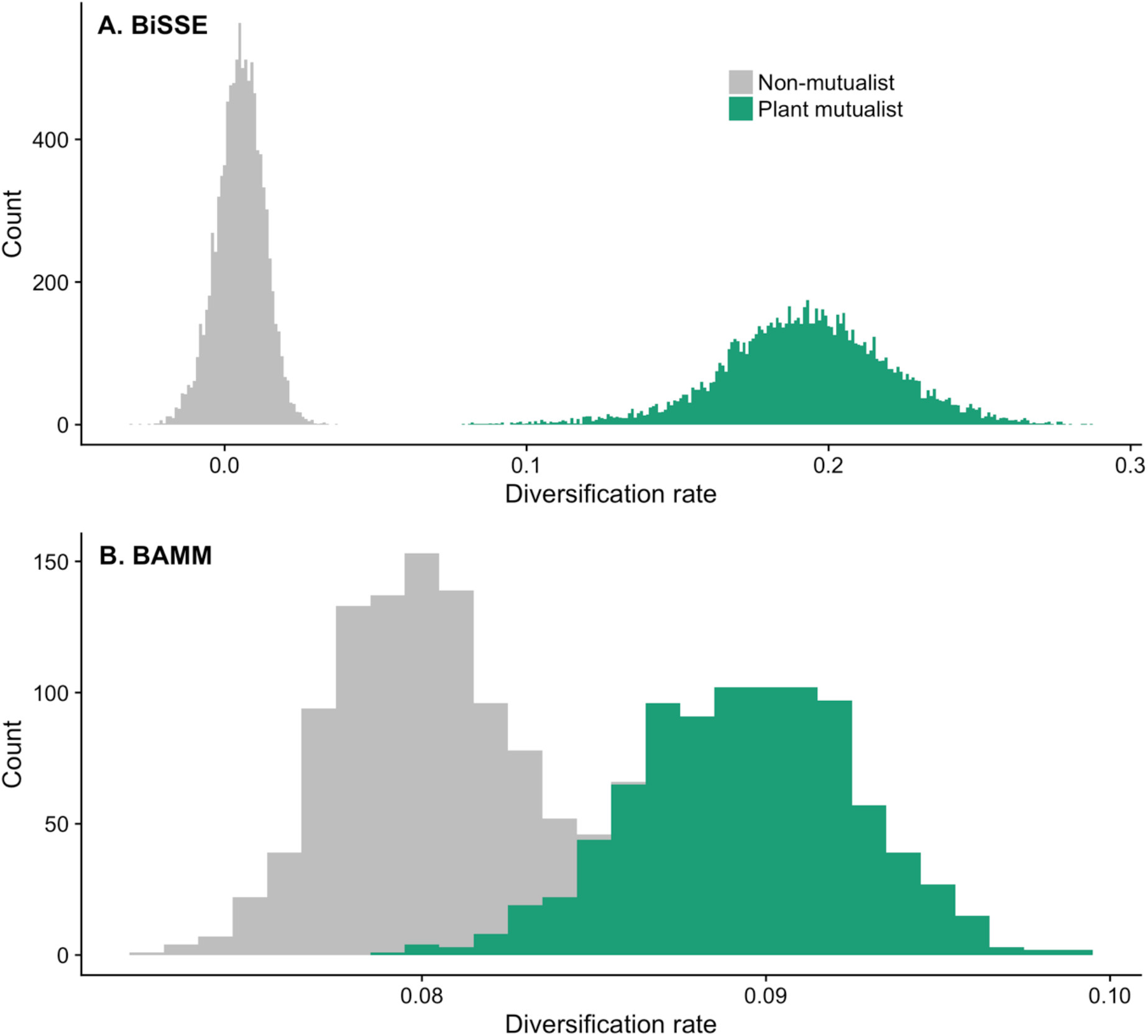
Diversification rates from (A) BiSSE and (B) BAMM analyses for ant lineages that are plant mutualists (green) and ant lineages that are not plant mutualists (grey).

**Table 1.** Parameter estimate summary from MCMC for BiSSE analysis and post-burn-in MCMC result summary for BAMM analysis. The mean and (standard deviation) of parameters are reported.

|  | BiSSE | BAMM |
| --- | --- | --- |
| **Speciation<sub>0</sub>** | 0.11 (0.017) | 0.13 (0.0073) |
| **Speciation<sub>1</sub>** | 0.69 (0.13) | 0.15 (0.010) |
| **Extinction<sub>0</sub>** | 0.10 (0.019) | 0.045 (0.0082) |
| **Extinction<sub>1</sub>** | 0.50 (0.16) | 0.064 (0.0011) |
| **Q<sub>01</sub>** | 0.017 (0.0039) | - |
| **Q<sub>10</sub>** | 0.20 (0.028) | - |
| **Diversification<sub>0</sub>** | 0.0052 (0.0078) | 0.080 (0.0022) |
| **Diversification<sub>1</sub>** | 0.19 (0.028) | 0.089 (0.0032) |
| **Overall Rate<sub>1-0</sub>** | 0.19 (0.033) | 0.0098 (0.0023) |

## Discussion

Our results build on previous research^21,22^ showing how automated methods can reliably and efficiently assemble large trait databases for answering macroevolutionary questions. Text-mining generated trait data for almost twice the number of ant species as in the most comprehensive phylogeny available (Nelson & Moreau, unpublished data), and we had overlapping trait data and phylogenetic information for 795 ant species. We used these data to test whether plant-ant lineages have elevated diversification rates, a hypothesis that was supported by both BiSSE and BAMM models. Previous research has similarly found that partnering with ants for defense or seed dispersal accelerates plant diversification^3,4^, suggesting that ant and plant lineages are either responding to the same external factor(s) affecting diversification (e.g., biogeography), or that ants and plants cooperate-and-radiate. Thus, in addition to having direct effects on both ant and plant partners, mutualism evolution may have long-term implications for lineage diversification, such that mutualism results in reciprocal diversification in interacting lineages.

Text-mining was a successful and efficient method for assembling a large trait database, and was primarily limited by the availability of abstracts. The species accumulation curves (Figure 2) suggest that with an even larger corpus, we could acquire trait data for many more ant taxa, and especially that we would identify many more EFN-visiting and seed-dispersing ant species. Our text-mining extracted similar trait information as what is normally assembled manually, and more laboriously, from the primary literature; an automated approach could also vastly improve datasets for meta-analyses, ecological network analysis, etc. Although our text-mining algorithm returned a small number of false positives because we took any co-occurrence of a trait term and an ant species name in an abstract as evidence of an association, the number of false positives declined rapidly as trait terms and ant names were found together in more abstracts (Figure 3), again suggesting that a larger corpus would help to reduce noise in the text-mining output. We also text-mined only abstracts; assuming abstracts of papers on ant-plant mutualisms are less likely than main texts to mention non-mutualistic ants in passing, our method should be more conservative regarding false positives, but of course full-text articles contain more trait information. To improve on our text-mining method, considering the proximity between words or the frequency of words, or a more restrictive rule for how often an ant name and trait term need to co-occur, could help to further reduce the frequency of false positives. However, a more restrictive rule may not be necessary with a sufficiently large corpus, as our comparison between the text-mined and hand-compiled data sets shows that most of the seed-dispersing ants missed by the text-mining were described in papers not in our corpus.

Automated downloading of large numbers of abstracts proved difficult as most are behind paywalls, and even with institutional subscriptions, it is challenging to download publications en masse^26^. Thus, more open access publications could increase the efficacy and benefits of automated data collection methods such as text-or data-mining. Nonetheless, using text-mining to extract data from published papers could be applied to a variety of research questions, given that data is readily available in word combinations.

Overall, the evolution of ant-plant mutualisms was associated with higher ant lineage diversification; we found a positive effect of mutualism evolution on ant diversification in both analyses, although the effect size was smaller in the BAMM than in the BiSSE analysis (Table 1, Figure 4). We report both speciation and extinction values from the BiSSE model but focus our discussion on diversification rate estimates. In the BiSSE analyses, although the traits evolve independently multiple times, mutualism is very frequently lost (Table 1). One possibility is that mutualism evolves, the ant lineage radiates, and then mutualism is secondarily lost multiple times.

This would explain the effect of mutualistic traits on diversification, despite the high mutualism loss rate. Perhaps a more likely explanation is that our trait dataset may include a high number of false negatives that BiSSE models reconstruct as secondary losses. If an ant name and trait term co-occur in enough abstracts, there is little doubt that the ant is a plant mutualist, but it is more challenging to be sure that ants named in abstracts that do not also contain trait terms are truly non-mutualistic.

Ant-plant interactions are often diffuse, generalized mutualisms^27^ and theory predicts that mutualistic lineages are prone to extinction^6,7^. For example, in angiosperms, domatia are thought to be an evolutionary dead end; they appear to go extinct easily in unfavourable conditions, or are as easily lost as they are gained^13^. However, although ants appear to revert easily back to a non-mutualistic state, we still found that mutualism positively affects ant diversification (see Figure S2 for ancestral state reconstruction). Our results also indicate that there are elevated extinction rates in mutualistic ants but the effect of this on diversification is swamped by the positive effect of mutualism on ant speciation. Compared to seed-dispersing ants, ants that nest in domatia or consume EFN or food bodies confer very different benefits to plants (dispersal versus defense, respectively) but most of these ants nonetheless receive similar rewards—mainly food, but also housing for domatia-nesting ants. These rewards could expand the geographic range or realized niche of an ant lineage. Similar arguments have been made to explain how the evolution of EFNs or elaiosomes has accelerated plant diversification^34^. In combination with previous work on angiosperm diversification, our finding that ant-plant mutualisms also spur ant diversification suggests ants and plants have diversified in tandem via “cooperate-and-radiate” co-evolution.

## Methods

### Sources of phylogenetic information

We used a phylogenetic tree that includes 1,731 ant species provided to us by Drs. Matthew Nelsen and Corrie Moreau, applying the drop.tip function in the R^28^ package *ape^29^* to include only tips for which we had trait data. The tree is a compilation of previously published ant trees and provides the most up to date inference of evolutionary relationships among ants. However, since there are 14,416 recognized valid ant species names (B. Fisher, personal communication), even a tree with 1,731 ant species is woefully under sampled. The recent phylogenetic comparative methods we used account for incomplete taxon sampling, but these methods are not without error^25^. We did not account for phylogenetic uncertainty, as such analyses are computationally intensive for many phylogenetic comparative methods.

### Sources of trait data

We text-mined 89,495 titles and abstracts from two sources, after removing duplicates prior to analysis: 1) 62,988 abstracts from *FORMIS:* A Master Bibliography of Ant Literature, containing all known ant literature through to 1996^30^, and 2) 52,885 ant-related publications available through Springer’s Application Portal Interface (API); Springer publishes numerous journals with substantial ant-related content, including *Oecologia, Insectes Sociaux, Arthropod-Plant Interactions*, and others. We used Springer’s API to search titles and abstracts for ant species names and plant traits that facilitate ant-plant mutualisms (Table S1). We downloaded the abstracts of the Springer articles that mentioned the ant species or trait terms on our list. One of us (CGL) provided a hand-compiled list of seed-dispersing ants from reading 180 journal articles; we supplemented the text-mining results with data from this hand-compiled dataset.

### Text-mining approach

Because a significant amount of information can be discerned from word combinations alone^31^, text-mining can be an effective tool to extract information from the published scientific literature. We took a text-mining approach to compile trait data on ant-plant associations; specifically, to determine which ant lineages consume food bodies, nest in domatia, visit EFNs, and disperse seeds. We created a term-document matrix to hold the trait data using the Pandas and NumPy packages in Python 2.7.12^32,33^. Our Python script used an *n*-gram approach that allowed us to identify short sequences of words, such as ant binomial nomenclature names or trait terms^31^. Our n-grams were: 1) all 14,416 currently valid ant species names in a global list of ants, excluding extinct taxa, and 2) a list of traits related to ant-plant mutualisms (Table S2). The resulting term-document matrix identifies whether 1) each ant name appeared without a trait or 2) each ant name appeared with a trait, in each publication’s title or abstract. We text-mined trait terms in four broad categories (domatia, extrafloral nectar, food bodies, and seed dispersal) to capture many different types of ant-plant mutualisms. We used the term-document matrix to score each mutualism category as a discrete binary trait for each ant species, and then further combined all the data into a single binary “plant mutualist” category.

### Text-mining validation

If an ant species name never occurred in the same abstract as a trait term (Table S2), but appeared in at least one abstract without a trait term, it was scored as an absence (0). If an ant species name co-occurred with a trait term in at least one abstract, it was scored as a presence (1). Both false positives and false negatives are a concern with text-mined trait data, although it is worth noting that other methods of gathering trait data can also be error-prone at such a large scale. Nonetheless, we validated our text-mined trait data in several ways. First, we scrutinized cases in which ant species names co-occurred with trait terms in 5 or fewer abstracts. We read each of these abstracts and manually scored the ant species in question as a true (1) or false (0) association with the trait term. We compared these manual scores to the text-mining results to determine how the number of false positives declined as the number of abstracts containing ant names and trait terms increased. The trait data for the manually checked abstracts were corrected if required for all subsequent analyses. Although we removed most duplicate abstracts from our corpus during initial processing, because not all abstracts were identically formatted, we discovered a few additional duplicates while reading abstracts and adjusted the dataset accordingly. We also calculated species accumulation curves, in which we determined how the number of unique plant-ant and non-plant-ant species increased as a function of the number of abstracts that were text-mined. Finally, we also compared the text-mined data to our hand-compiled data set.

### Lineage diversification analyses

Rabosky and Goldberg^34^ showed that low transition rates and rare shifts can cause high type 1 error in SSE models, but this is not a problem in our analyses because ant-plant mutualisms have evolved many times independently across the ant phylogeny (Figure 1). However, Rabosky and Goldberg^34^ also highlighted a SSE model inadequacy in which neutral traits spuriously influence diversification rates. For this reason, we assessed the influence of being a plant-mutualist on lineage diversification in two ways, using: 1) Binary State Speciation and Extinction^24^ (BiSSE) models implemented using the *diversitree^35^* package in R^28^ and 2) Bayesian Analysis of Macroevolutionary Mixtures (BAMM) models using bamm 2.5.0 and *BAMMtools*^36^ in R^28^.

BiSSE estimates the transition rate between states (q_01_ and q_10_) as well as state-specific extinction and speciation rates (mu_0_ and mu_1_ and lambda_0_ and lambda_1_, respectively). We ran a BiSSE model using a global proportion of missing data, specifically the number of species in the tree divided by the total number of currently recognized ant species. To test whether mutualist and non-mutualist lineages have different diversification rates, we ran a Markov Chain Monte Carlo (MCMC) BiSSE analysis with an exponential prior with rate 1/2r, where r is the independent diversification rate of the character. An initial MCMC was run with a tuning parameter of 0.1 for 1,000 generations. The revised tuning was calculated from the width of the middle 90% of the posterior samples from these initial runs. The MCMC analysis was subsequently run for 10,000 generations. We estimated net diversification (speciation - extinction) rates in mutualistic and non-mutualstic ant lineages. We calculated 95% credible intervals of the posterior samples for each parameter from the BiSSE run.

We also ran BAMM analyses three times on the same tree. The BAMM was run for 10 million MCMC generations, sampling the parameters after every 100,000 generations. We included an overall proportion of ant species in the tree over the total number of indentified ant species in order to indicate how much clade information was missing to account for incomplete taxon sampling (see supplementary file). Rate priors were calculated for each tree using *BAMMtools*^36^ and convergence of the BAMM runs was also tested using the R^28^ package *coda^37^*. To assess if diversification rates differed in mutualistic and non-mutualistic ant lineages, we used the subtreeBAMM and getcladerate functions in *BAMMtools*^36^ to assess whether there were areas of the tree in the mutualist state that had higher diversification rates than the non-mutualist state. We used the BAMM output to assess the diversification rate of mutualistic and non-mutualistic ant lineages.

## Data availability

Trait and species information used in the text-mining will be available in DataDryad upon acceptance.

## Code availability

Text-mining code (Python script) and lineage diversification code (R scripts for BiSSE and BAMM analyses) will be available in DataDryad upon acceptance. Scripts are also available on github (respository name: Cooperate-and-Radiate).

## Acknowledgements

We thank Matthew Nelsen, Corrie Moreau, and Brian Fisher for providing us with phylogenetic and taxonomic information. We also thank Luke Mahler, James Thomson, Santiago Claramunt, Asher Cutter, Marjorie Weber, Rebecca Batstone, Emily Dutton, Jason Laurich, Anna O’Brien, Shannon-Meadley Dunphy, Mitchel Trychta, and Tia Harrison for their help and feedback over the course of this project. We acknowledge the financial support of a Natural Sciences and Engineering Research Council (NSERC) Discovery Grant and Discovery Accelerator Supplement to MEF.

## Author contributions

MEF, PJM, and KMK conceived the study and approach. PJM and KMK obtained abstracts, KMK and ES wrote and edited the text-mining script. CGL generated the hand-compiled list of papers.

KMK ran phylogenetic analyses and lead the writing of the manuscript, with contributions from MEF.

## Additional information

Supplementary information is available for this paper.

Correspondence and requests for materials should be addressed to KMK.

